# The Block Object Storage Service (bossDB): A Cloud-Native Approach for Petascale Neuroscience Discovery

**DOI:** 10.1101/217745

**Authors:** Robert Hider, Dean M. Kleissas, Derek Pryor, Timothy Gion, Luis Rodriguez, Jordan Matelsky, William Gray-Roncal, Brock Wester

## Abstract

Large volumetric neuroimaging datasets have grown in size over the past ten years from gigabytes to terabytes, with petascale data becoming available and more common over the next few years. Current approaches to store and analyze these emerging datasets are insufficient in their ability to scale in both cost-effectiveness and performance. Additionally, enabling large-scale processing and annotation is critical as these data grow too large for manual inspection. We provide a new cloud-native managed service for large and multi-modal experiments, with support for data ingest, storage, visualization, and sharing through a RESTful Application Programming Interface (API) and web-based user interface. Our project is open source and can be easily and cost-effectively used for a variety of modalities and applications.

## 1 Introduction

Mapping the brain to better understand cognitive processes and the biological basis for disease is a fundamental challenge of the 21st century that is only now emerging as a realistic endeavor, realizing the dreams of early neuroscientists such as Ramón y Cajal, who were limited to sketching brain maps in ink, one neuron at a time. Technological advances in neuroscience have exploded over the last ten years, making it almost routine to image high-resolution (sub-micron) brain volumes in many laboratories around the world [1, 2, 3, 4, 5].

As new tools for interrogating tissue at these resolutions advance and become more common, a centralized data archive is needed to enable the large (tera- to petascale) storage, visualization, and discovery processes, and address some of the challenges identified by the community [3, 6]. While research groups are beginning to embrace data archives, most treat the system as simply a place to deposit finalized data, with raw datasets generated and stored in a custom format and analyzed and inspected with custom software. At this scale, it is quickly becoming impossible for researchers to characterize many of the underlying properties; indeed, for petascale volumes, it is likely that most of a data volume is never viewed in detail by a human. Additionally, conventional approaches for automatically or semi-automatically reconstructing neuronal maps focus on building methods for small volumes, and scaling these tools to operate on multi-terabyte or petabyte data volumes is often either unachievable or significantly beyond the capabilities and budgets of a research group.

Large datasets are a resource incredibly rich in scientific content, which should be shared with others to best leverage the investment of time and resources and to fully exploit the value of the data. Due to the challenges in collection, storage, and analysis of terascale and petascale data volumes, few public datasets of this size are routinely shared, although many such volumes exist on local storage and a deluge of new data is forthcoming [7, 8, 9].

We considered use cases such as the first fully-automated pipelines for processing and assessing X-ray Microtomography (XRM) [10] and electron microscopy (EM) datasets [1, 2, 4] and work by many academic laboratories around the world to understand state-of-the-art approaches and their limitations, and emphasize that high-performance and scalable data storage is an essential component of any modern connectomics effort. In designing our Block and Object Storage Service Database (`bossDB`), we researched several related efforts, including DVID [11] which excels in versioned terascale storage and CATMAID [12] which provides a mature manual annotation platform. We previously worked with NeuroData to develop ndstore [13], which originated and implemented many of the design principles necessary to store and access high-dimensional imaging datasets, including an efficient internal data representation and associated spatial indexing scheme; the Reusable Annotation Markup for Open Neuroscience (RAMON), an annotation schema for connectomics [14]; an API to remotely access services; and MATLAB and Python toolkits to facilitate usability. Based on this prior research and an understanding of the evolving requirements driven by new and maturing imaging modalities, we created a robust, cloud-native petascale datastore with a number of services and support tools (Figure 1).

**Figure 1:**
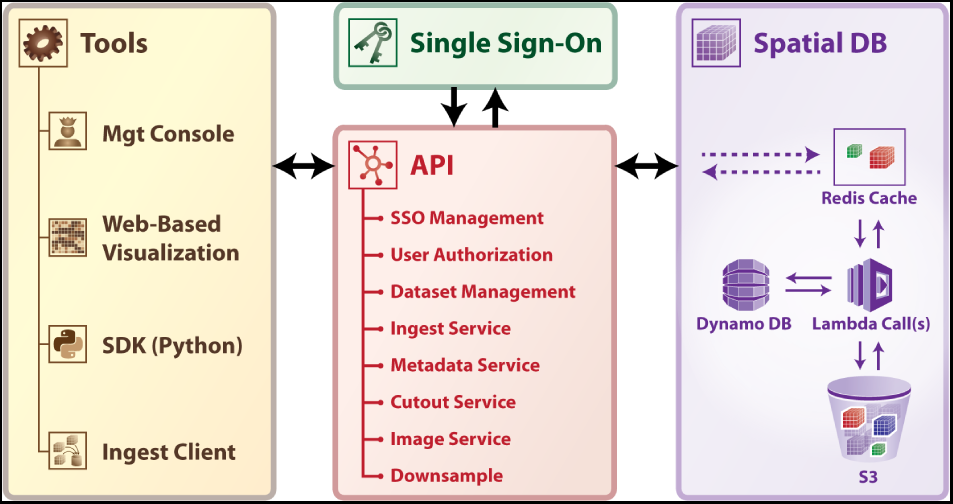
A high-level schematic of `bossDB` platform.

## 2 Methods

To enable large-scale, collaborative research we developed and deployed a cloud-native data archive to support the storage, analysis, and sharing of large spatial datasets. Service-oriented architectures have continued to grow in popularity and possess many appealing properties when designing a cloud-based data archive [15]. Our solution, called the Block and Object Storage Service (`bossDB`), is currently deployed in the Amazon Web Services (AWS) cloud and has been robustly architected to leverage cloud capabilities and ensure a highly-available, scalable, and cost-efficient system.

### 2.1 Spatial Database

The spatial database is the foundation of `bossDB`, and uses the strengths of the cloud to efficiently store and index massive multi-dimensional image and annotation datasets (i.e. multi-channel 3D image volumes). A core concept is our managed storage hierarchy, which automatically migrates data between affordable, durable object storage (i.e. Amazon S3) and an in-memory data store (i.e. Redis), which operates as read and write cache database for faster IO performance. This allows for storage of large volumes at a low cost, while providing low latency to commonly accessed regions. We utilize AWS Lambda to perform parallel IO operations between the object store layer and memory cache layer and DynamoDB for indexing. These serverless technologies allow `bossDB` to rapidly and automatically scale resources during periods of heavy operation without incurring additional costs while idle.

The `bossDB` spatial database is designed to store petascale, multi-dimensional image data (i.e. multichannel 3-dimensional image volumes, with optional time series support, Figure 2, 3) and associated co-registered voxel annotations. In this context, voxel annotations are unsigned 64-bit integer labels stored in a separate channel that is in the same coordinate frame as the source image data. Each unique uint64 value represents a unique object (e.g., neuron, synapse, organelle). A user can then label voxels that have some semantic meaning, typically the result of manual annotation or automated processing. The database maintains an index of annotation locations, enabling efficient spatial querying and data retrieval (Figure 4).

**Figure 2:**
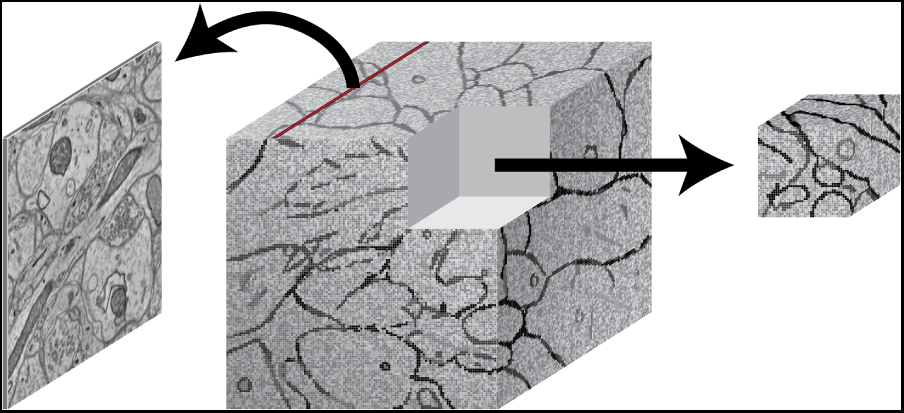
An illustration showing image slices (left) being composed into 3D cuboid volumes (middle). Arbitrary requests may be made to extract image regions of interest (right).

**Figure 3:**
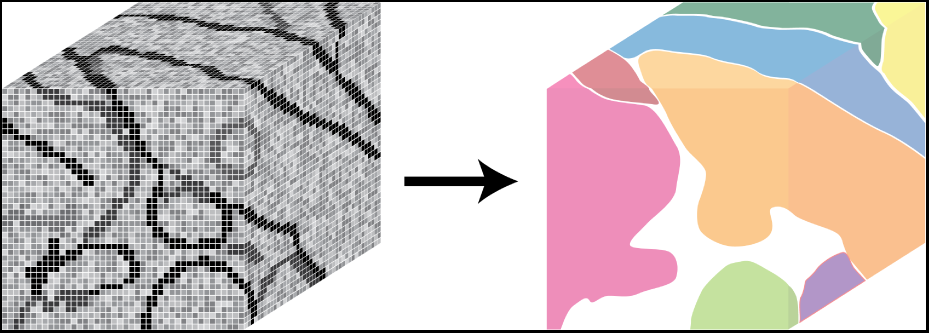
An illustration showing annotations, composed of voxel labels (left) and how a unique annotation identifier can represent a unique object in the image data (right).

**Figure 4:**
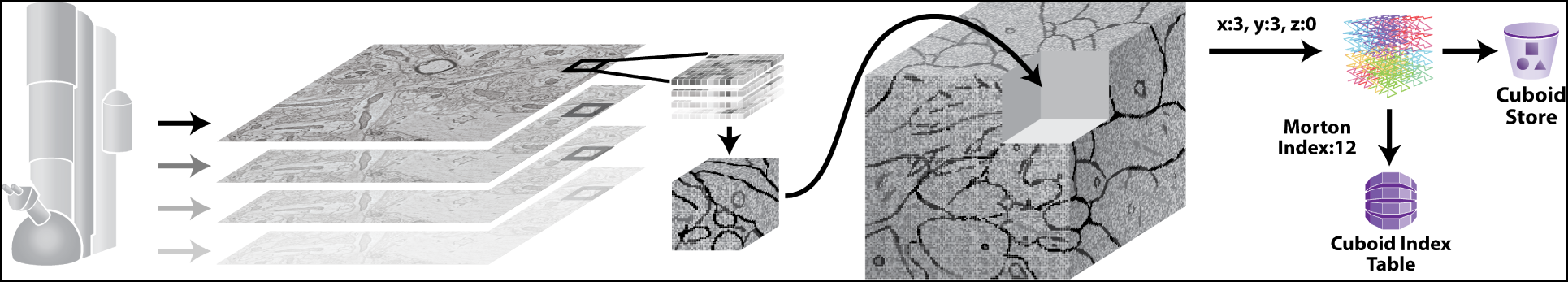
An illustration showing how large 2D image slices generated by an electron microscope are reformatted as cuboids, which fit into a larger 3D volume, indexed using a z-order curve.

The internal representation of volumetric data is inherited from previous collaborative work with the NeuroData ndstore project [13, 16]. Here, we operate on small cuboids, or 3D chunks of data (e.g., 512 × 512 × 16 voxels), which are stored in Amazon S3 as compressed C-order arrays. Cuboids are indexed using a Morton-order space-filling curve, which maps the 3D location of each cuboid to a single dimension. In addition, annotations are indexed so `bossDB` can quickly retrieve which annotation IDs exist in an individual cuboid, and in which cuboids a unique ID exists. With these indices, all of which are stored in auto-scaling Amazon DynamoDB tables, the `bossDB` API can provide spatial querying of annotations by ID and efficient retrieval of arbitrary data volumes. The database will also render and store a resolution hierarchy through down sampling of a dataset, which is critical for visualization applications to efficiently provide low-resolution views and useful when processing large datasets.

The spatial database supports various bit-depths (i.e. uint8, uint16 image channels and uint64 annotation channels) and we will provide additional bit-depth and data formats as needed.

### 2.2 Single Sign-On (SSO) Identity Provider

A centralized and standalone authentication server provides single sign-on functionality for `bossDB` and integrated tools and applications. This allows `bossDB` to control permissions internally and operate securely, while maintaining the ability to federate with other data archives in the future.

A robust authentication and authorization system provides many important benefits, such as the ability to keep some data private while other data public and to control what actions individual users are permitted to perform. We use the open source software package Keycloak as an identity provider to manage users and roles. We created a Django OpenID Connect plugin to simplify the integration of services with the SSO provider.

Our identity provider server intentionally runs independently from the rest of `bossDB` system, forcing the `bossDB` API to authenticate just like any other SSO integrated tool or application, and making future federation with other data archives or authentication systems easy. The Keycloak server is deployed in an auto-scaling group that sits behind an Elastic Load Balancer.

### 2.3 Application Programming Interface (API)

As the primary interface to `bossDB`, the API provides a collection of versioned, RESTful web services. It enforces access permissions and organizes data in a logical data model for spatial and functional results. Because the API is versioned, the `bossDB` storage engine can support significant changes while still maintaining backwards compatibility with legacy applications and tools. All requests to the API are authenticated through the SSO service or via a long-lived API token, which enables tracking usage and throttling requests as needed to manage cost and ensure reliable performance (e.g., high bandwidth power user vs. a limited guest user). The services `bossDB` provides are summarized below:

#### 2.3.1 SSO Management and User Authorization

A set of services to manage users, roles, groups, and permissions. Roles limit what actions a user can perform on the system, while permissions limit what data users can access or manipulate. Permissions are applied to `bossDB` datasets via groups, making it easy to manage and control access for both individuals and teams. Through the application of permissions, a researcher or administrator can choose to keep a dataset private, share with collaborators, or make it publicly available.

#### 2.3.2 Dataset Management

The `bossDB` API organizes data into a logical hierarchy to group related data together (e.g., source image data and associated annotations, 2-photon and EM datasets from the same tissue sample). This service provides interfaces to create and manage datasets and their properties.

#### 2.3.3 Ingest

A critical challenge when using a centralized data archive is the ingest of large datasets, as users will always locally organize and store data they generate in unique ways. The Ingest Service facilitates moving large datasets from local storage into `bossDB` by decoupling the upload of data to the cloud and ingesting of data into the spatial database, allowing independent scaling and failure recovery. The service provides methods to create a new ingest job, monitor the status of a job, join an upload client worker to a job, and cancel a job.

##### 2.3.3.1 Tile ingest

As demonstrated in Figure 5, the ingest process directly leverages AWS infrastructure, massively scaling on demand. First, using the ingest client a user uploads an ingest job configuration file to the API (1) which populates a task queue, enumerating all tiles that must be uploaded, and returns temporary AWS credentials. Next, the ingest client retrieves a task from the Upload Task Queue (2), and loads the requested local file into memory as an image tile (3) and uploads the tile data to an S3 bucket (4). The ingest client then writes a message to the index queue signaling it is finished with this tile (5). An AWS Lambda automatically fires when a message enters the Index Queue and it uses DynamoDB to track which tiles are successfully written to the tile bucket (6)(7) and when enough tiles in a region have arrived to generate the `bossDB` cuboid data representation, a second Lambda function is triggered (8). This Ingest Lambda function then loads the specified tiles, reformats them into cuboids, inserts them into the Spatial DB S3 bucket, updates the Spatial DB cuboid index, and finally marks the temporary tiles for deletion (9). The ingest client supports both parallel and distributed operation, allowing users to maximize their network bandwidth.

**Figure 5:**
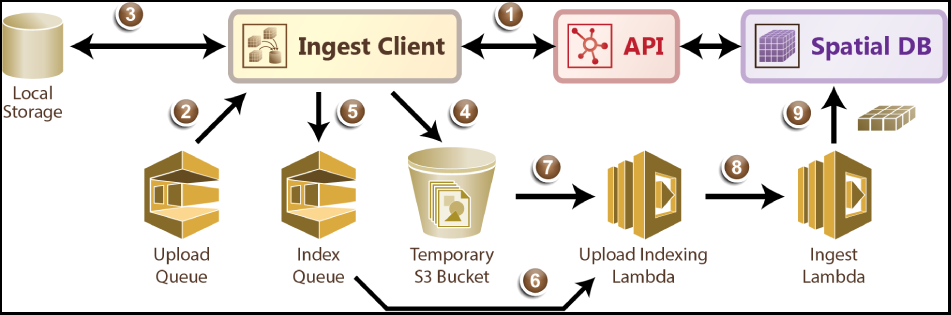
A diagram outlining the ingest process.

##### 2.3.3.2 Volumetric ingest

The ingest process also supports uploading threedimensional chunks of data in the CloudVolume format [17]; this interface can be straightforwardly extended to other formats. Similar to Tile Ingest, the ingest-client is used to upload an ingest-job configuration file to the API, populating a task queue with all chunks to be uploaded. The ingest client then retrieves a task from the Upload Task Queue, and loads that chunk into memory. The memory chunk is divided into multiple `bossDB` cuboids (512 x 512 x 16) and each cuboid is uploaded to an AWS S3 bucket. Upon uploading, the S3 update will trigger an AWS Lambda that copies the cuboid into main s3 store, adds an entry in DynamoDB and marks the original cuboid for deletion.

#### 2.3.4 Dataset Metadata

`bossDB` can store arbitrary key-value pairs linked to data model items, which is useful to track experimental metadata and provenance (e.g., voxel size, animal information, annotation algorithm used). This service provides an interface to query, create, update, and delete key-value pairs associated with a dataset.

#### 2.3.5 Cutout

A service to interact with the Spatial Database by reading and writing arbitrary data volumes. While `bossDB` stores all data internally using a standardized format, the cutout service uses HTTP content negotiation to determine the data format of a request, allowing `bossDB` to easily support user-specified formats when uploading or downloading data (e.g., compressed C-order blob, hdf5 file, pickled numpy array). This service enables scalable analytics by letting users access arbitrary chunks of data in parallel, perform automated processing, and write the annotation result back to `bossDB`. It also supports querying for the spatial properties of annotations, such as the bounding box of an annotation or identifying which annotations exist within a region.

#### 2.3.6 Image

In addition to our volumetric cutout service, we provide an image service to meet common user needs, which retrieves a 2D slice of data from the spatial database, along one of the three orthogonal planes (i.e., XY, XZ, YZ), encoded as an image file. Again, HTTP content negotiation is used to determine the format of the response (e.g., png, jpeg). The service supports arbitrary image sizes or a fixed tile size, which is often used by visualization tools.

#### 2.3.7 Downsample

To allow users to quickly assess, process, and interact with their data, we need to iteratively build a resolution hierarchy for each dataset by downsampling the source data. This is a workflow that is run infrequently and on-demand, and needs to scale from gigabytes to petabytes of data. We developed a serverless architecture built on AWS Step Functions to manage failures and track process state. AWS Lambda is used to perform the underlying image processing in an embarrassingly parallel, scalable fashion. This approach allows us to minimize resource costs while scaling on-demand in a fully-automated paradigm. It is also possible to perform a partial downsample when only a portion of the original dataset has changed, saving the time and expense of re-running the process on the entire database.

### 2.4 User Tools

User facing tools are required to make a data archive truly useful, easy to use, and well documented. We currently offer a web-based management console, an ingest client, and a client-side Python module for programmatic interaction. We have also integrated 3rd-party web-based data visualization tools. While `bossDB` API provides a rich interface to interact with the system, user friendly tools built on top of the API are critical to increase utility and adoption by the community. We expect this tool library to grow as users build on the core `bossDB` technologies.

#### 2.4.1 Web-based Management Console

`bossDB` has a web interface that lets users perform common actions directly in their browser (e.g., create a dataset, monitor an ingest job, share a dataset with a user). This console is the primary interface for most users and will expose much of the API’s functionality through an intuitive graphical interface. From the console, a researcher is able to manage datasets, discover new data, and launch the visualization tool.

#### 2.4.2 Web-based Visualization

A critical capability to any data archive is the ability to easily visualize stored data. Whether inspecting ingested data, exploring a dataset, or sharing an interesting sample with a collaborator, often the most common interaction with stored data will be through visualization. We integrated a version of Neuroglancer [18] to let users visually explore data stored in `bossDB`.

#### 2.4.3 Ingest Client

We have developed an open source ingest client in Python to manage uploading data to `bossDB`. The ingest process operates on a upload task queue which contains tasks specifying individual 2D tiles or 3D chunks of data to upload. To deal with the unique formats and file organization methods of diverse users, the client uses a simple plug-in design to import custom snippets of code responsible for taking a task, finding the right file, and loading the data into memory, which is then uploaded by the client. The work queue design allows copies of the client to be run distributed across compute nodes and in parallel on a single machine, substantially increasing throughput.

#### 2.4.4 Python Software Development Kit (SDK)

To support developers and researchers who want to programmatically interact with `bossDB`, we provide a pip-installable Python library that abstracts much of the API’s complexity away from the user. Data cutouts of arbitrary size can be efficiently retrieved from our archive, enabling easy integration with analytics tools. The current SDK, called `intern`, will continue to be expanded and supported to accommodate updates and additions to the existing `bossDB` system and user requests.

## 3 Results

### 3.1 Motivating Application

Many of our design requirements were motivated by the Machine Intelligent from Cortical Networks (MICrONS) Program [7]. This effort seeks to rapidly advance machine learning capabilities by creating novel machine learning algorithms that use neurally-inspired architectures and mathematical abstractions of the representations, transformations, and learning rules employed by the brain. To guide the construction of these algorithms, the program centers around massive co-registered functional (e.g., two-photon calcium imaging) and structural (e.g., EM and XRM) neuroimaging experiments aimed at estimating the synapse-resolution connectome of a 1 *mm*^3^ volume of cortex, represented by 2 petabytes of data, and using that information to constrain machine learning architectures. Our goal was to organize, store, and support the analysis of these large functional and anatomical datasets while enabling public dissemination.

### 3.2 Deployment

We envision that this data archive will facilitate neuroscience inquiries through an extensible, scalable process, with a sample process outlined in Figure 6. Below we describe our results to date for the MICrONS program datasets. Currently we serve approximately 50 regular users and 2 PB of compressed image data in our hosted system. In addition `bossDB` has 13TB of data from other collaborators.

**Figure 6:**
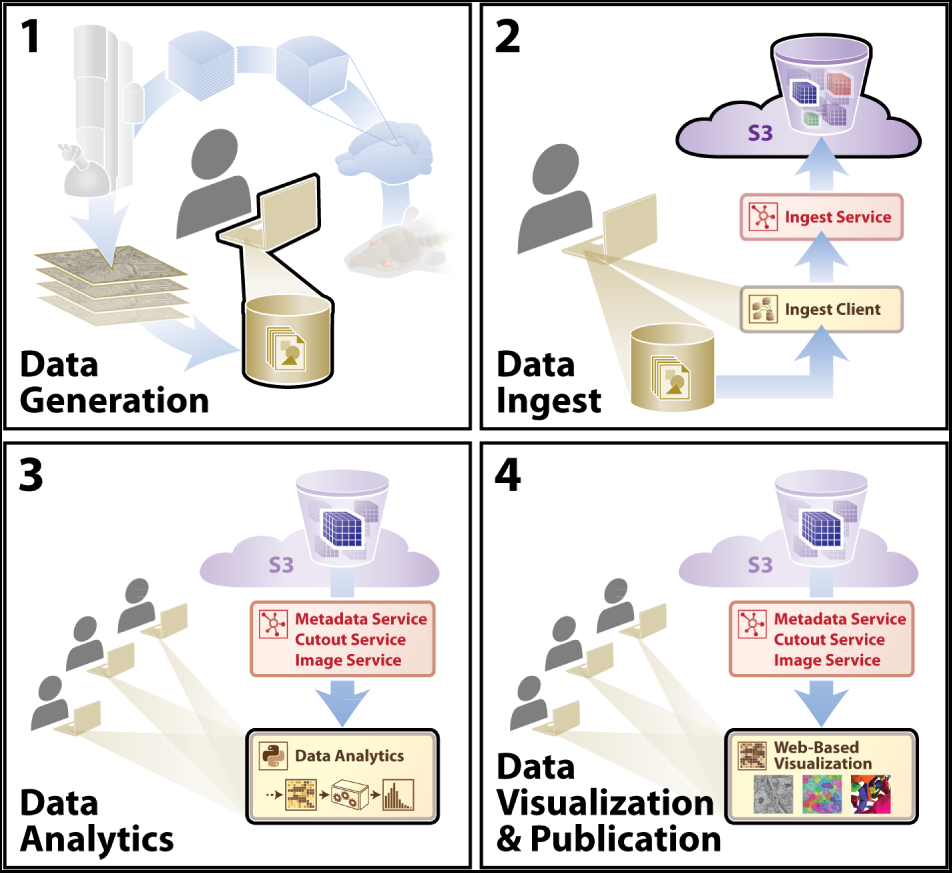
A diagram outlining an example user story showing utilization of the `bossDB` infrastructure. A typical research group collecting data for a hypothesis will move sequentially from (1)-(4). Other groups will extend these analyses using steps (3) and (4).

#### 3.2.1 Implementation

Figure 7 shows the architecture of `bossDB`. The system has two user facing services: Authentication and Web Server Endpoint, both of which sit behind AWS elastic load balancers. The system uses Keycloak servers in a high-availability configuration for single sign-on authentication. The web server endpoints use Django API, to provide access to the majority of the services in `bossDB`.

**Figure 7:**
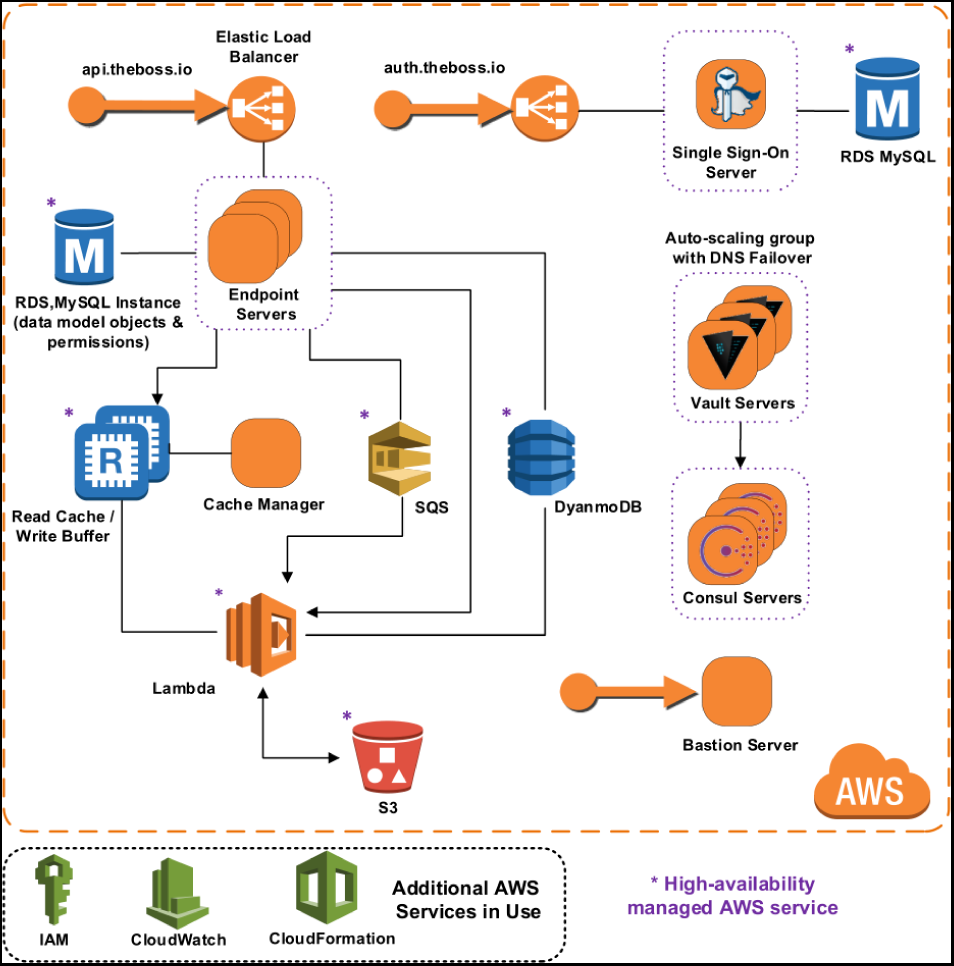
A high-level architecture diagram of `bossDB` as deployed using Amazon Web Services architecture.

`bossDB` uses serverless computing and storage, with AWS Lambda, SQS, S3 and DynamoDB to provide all of the other services mentioned in Section 2: Ingest, Metadata, Cutout, Image and Downsample. Using serverless computing and storage for these components will automatically scale with demand and eliminate the need to maintain components.

`bossDB` is installed using the AWS CloudFormation service along with Salt and Packer to manage our infrastructure. This allows us to quickly duplicate the environment for testing and development and even change instance sizes within the new environments.

#### 3.2.2 Data Generation

Researchers collect experimental data; stitching, alignment, and registration take part prior to upload to `bossDB`. Users create new resources in `bossDB` to identify and store their datasets, recording their experimental parameters and dataset properties (e.g., voxel dimensions, bit depth, spatial extent) prior to upload. An example screenshot from our web console is shown in Figure 8; this setup can be accomplished programmatically using `intern` as well.

**Figure 8:**
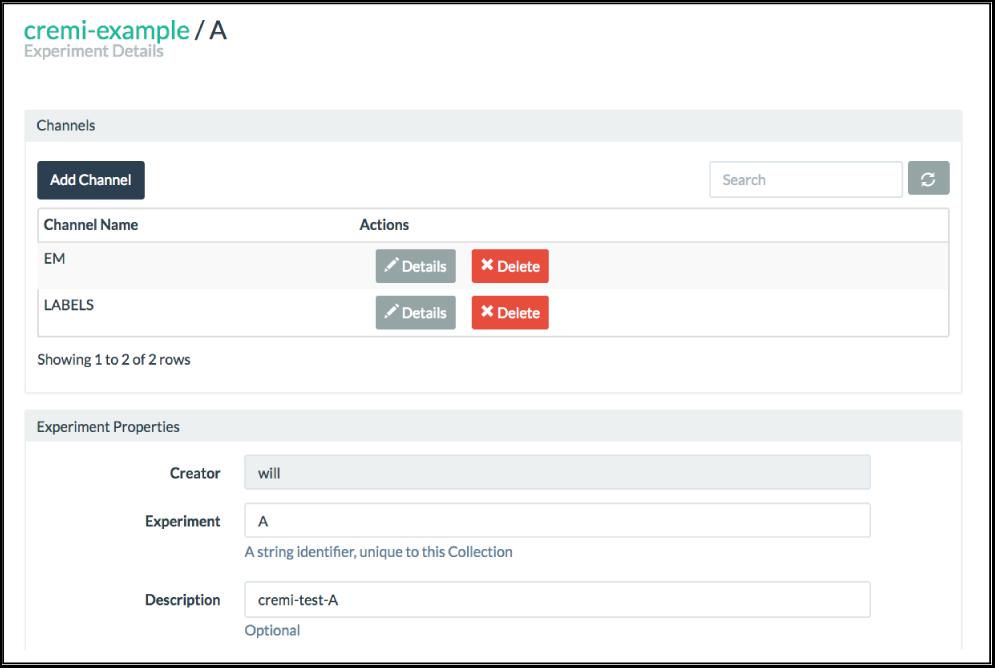
An example screenshot from our `bossDB` console.

#### 3.2.3 Data Ingest

Once available, a researcher uploads image data via one of several methods supported by `bossDB` (e.g., REST API, ingest client), safely and efficiently storing data in `bossDB`. Large datasets can be uploaded incrementally, with data available for read as soon as it has been ingested, providing access to collaborators in minutes, not months.

The ingest client has already been used to upload petabytes of EM and calcium imaging data; many of these uploads proceed without any intervention from the developer team with the system automatically scaling to meet user’s needs.

Recent testing of the ingest process reached a maximum sustained ingest throughput of 230 GB/Min (Figure 9) using the volumetric ingest-client into `bossDB`. The ingest client was run on 750 kubernetes pods across eight large servers uploading data from an AWS Bucket. AWS Lambda scaled to over 5000 concurrent executing functions to handle the load.

**Figure 9:**
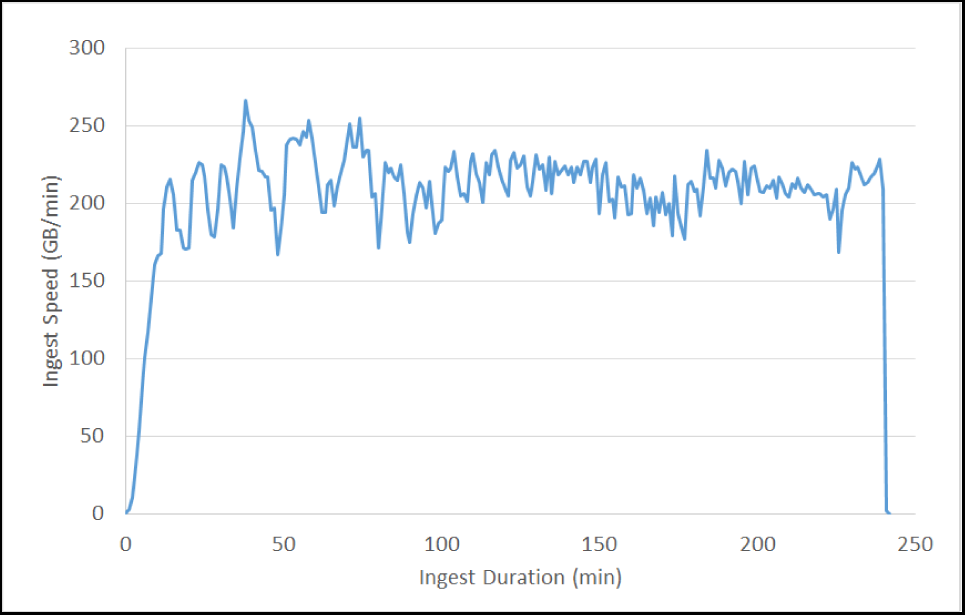
Volumetric Ingest throughput

To perform at this speed we were running 12 End-point servers sized with m4.2xlarge instances, a RDS database backed with a db.m4.xlarge instance, and DynamoDB table sized at 2000 read / 4000 write capacity.

This test shows the how `bossDB` will autoscale to meet demands (Figure 10). The same 3.2 million tiles were uploaded during each test. Each one used a different number of kubernetes pods running the ingest-client (100, 200, 400). `bossDB` automatically scaled endpoints, dyanmoDB read and write demand to handle the throughput efficiently.

**Figure 10:**
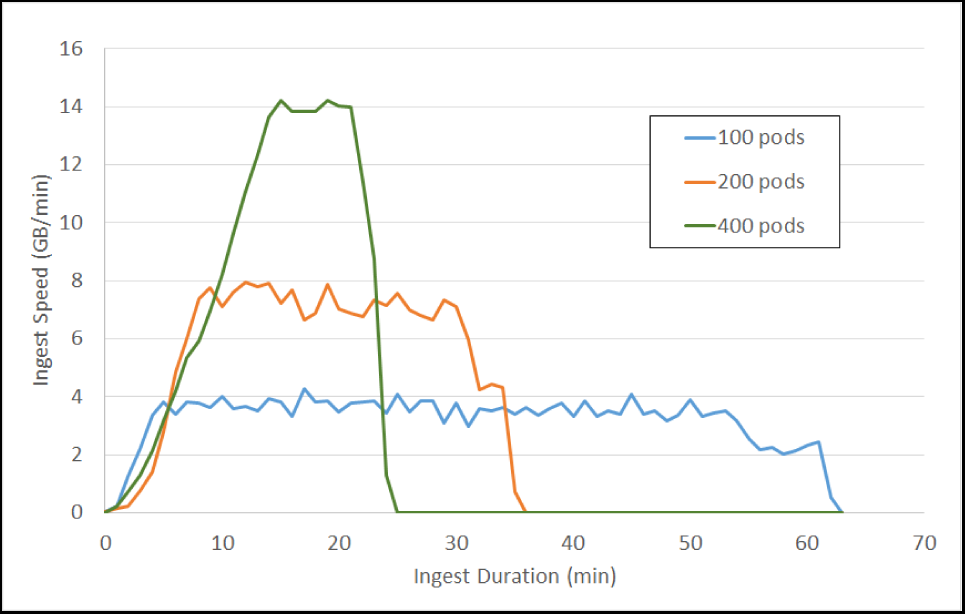
Tile Ingest throughput on demand

`bossDB` has monitoring capability at several levels. In Figure 11 you see a snippet of our Ingest Dashboard which allows the administrator to see how much stress any one component of the system is under. Notifications will also go out if any key components fail, and when the system hits cost milestones.

**Figure 11:**
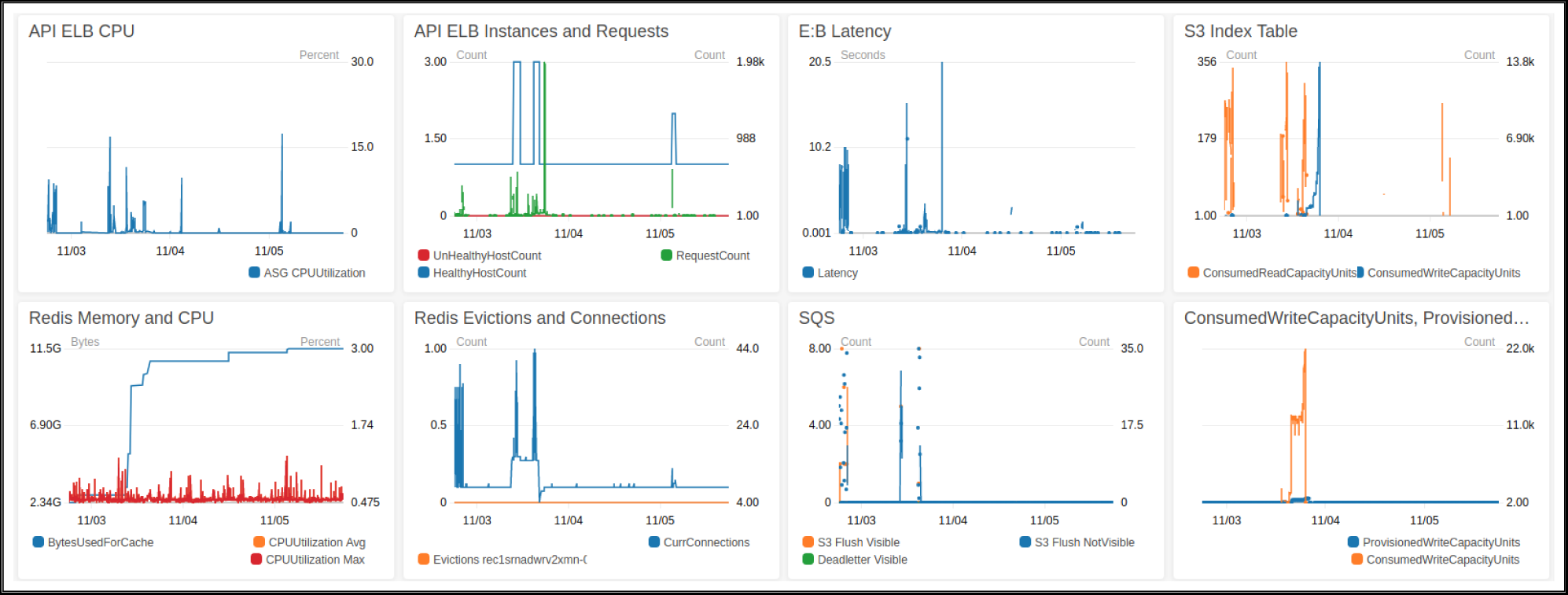
A CloudWatch dashboard monitoring during ingestion.

#### 3.2.4 Data Analytics

Many big data research analyses are enabled by `bossDB` features (e.g., standardized interfaces, arbitrary cutouts, spatial indexing), accelerating the scientific process.

One common use for `bossDB` is acting as a back-end for local data analysis pipelines. Users download chunks of data from `bossDB` using `intern` and process it to create annotation labels using humans or machines. The resulting annotation data is uploaded via a choice of methods (python API, ingest client), below we include an example of such use case.

~~~
% # import intern package
% **from intern**. remote. boss **import** BossRemote
% **import** numpy

% # initialize BossRemote
% rmt = BossRemote()

% # specify data location
% COLL_NAME = ‘test_collection’
% EXP_NAME = ‘test_experiment’
% CHAN_NAME = ‘test_channel’

% chan = rmt. get_channel(CHAN_NAME, COLL_NAME, EXP_NAME)

% # specify download coordinates and resolution
% x_rng = [0, 1024]
% y_rng = [0, 512]
% z_rng = [0, 10]
% res = 0

% # Download the cutout from the channel
% data = rmt. get_cutout(chan, res, x_rng, y_rng, z_rng)

# import intern package
**from intern**. remote. boss **import** BossRemote

# initialize BossRemote
rmt = BossRemote()

# specify data location
COLL_NAME = ‘test_collection’
EXP_NAME = ‘test_experiment’
CHAN_NAME = ‘test_channel’

# Create a reference to the channel resource: chan = rmt. get_channel(CHAN_NAME, COLL_NAME,
EXP_NAME)
# Or use a URL to identify the channel:
chan = f” bossdb://{ COLL_NAME}/{ EXP_NAME}/{ CHAN_NAME}”

# specify download coordinates and resolution x_rng = [0, 1024]
y_rng = [0, 512]
z_rng = [0, 10]
res = 0

# Download the cutout from the channel
data = rmt. get_cutout(chan, res, x_rng, y_rng, z_rng)
~~~

#### 3.2.5 Data Visualization and Publication

Data can be quickly visualized using applications such as NeuroGlancer (Figure 12).

**Figure 12:**
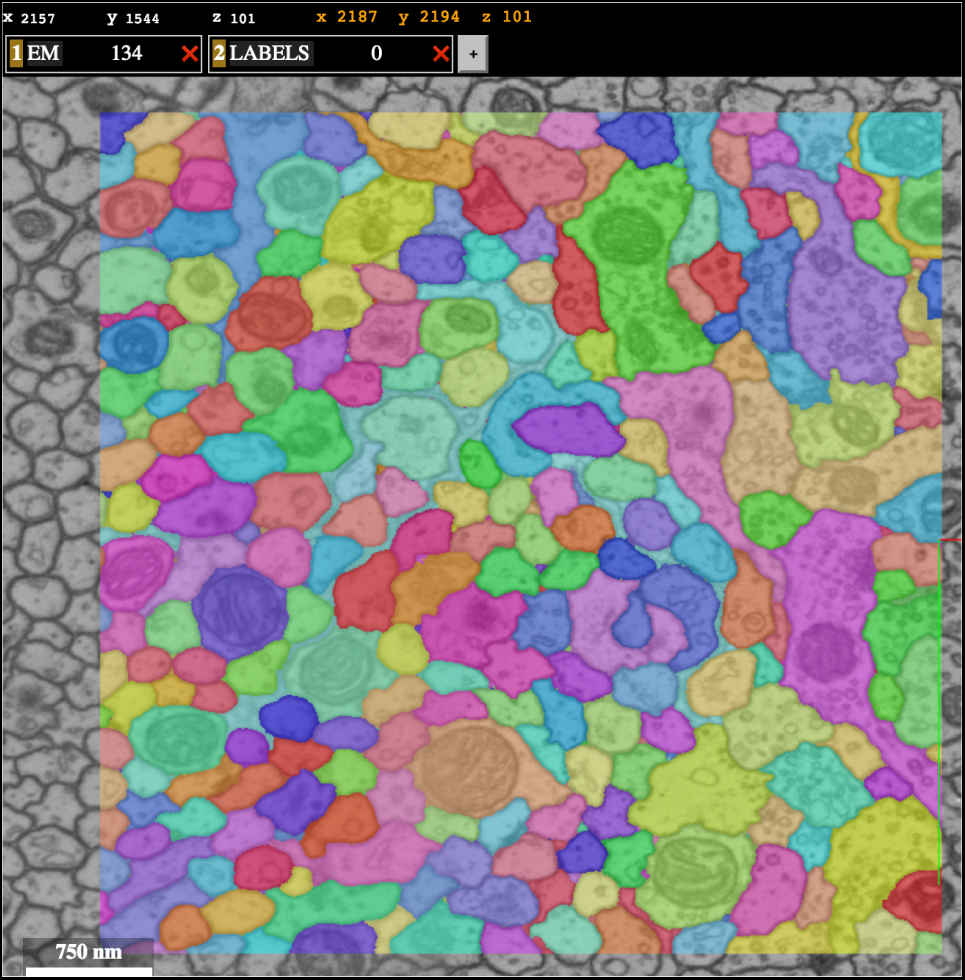
An example neuroglancer web visualization backended by `bossDB`, showing data from the recent CREMI challenge (www.cremi.org).

Data are published along with initial analysis, and made widely accessible through `bossDB`. Other research teams conduct additional analysis, extending and validating the existing scientific findings. By providing a standardized data format, it is easy for researchers to apply new techniques and test additional hypotheses.

## 4 Discussion

Our data archive will enable scientists to easily access and process large datasets, and to scale up their approaches with minimal alterations and without needing large local storage. Because the results are anchored to a universally-accessible datastore, it is easier for others to inspect the results, improve upon them, and reproduce processing pipelines by leveraging common interfaces.

When considering a cloud-native approach, vendor lock-in is one potential concern – as we not only use the AWS cloud to deploy `bossDB`, but have integrated many of its services into the system to substantially accelerate development and performance. To minimize the development impact of expanding to an additional cloud provider or on-premise cluster, future work is needed to create a layer of abstraction between the core software and AWS services. We plan to continue to develop towards a microservices style architecture, which will decrease coupling between sub-components. This will allow `bossDB` to be able to independently scale sub-components and increase the ability to easily deploy, update, and manage services. We believe that storage engines will continue to specialize around datatypes (e.g., multidimensional image data, video data, gene sequence data) and be applicable to multiple research communities through the creation of domain-specific APIs that maintain the unique formats, organization, and needs of that community.

We expect that as the community uses our data archive, additional tools will be developed to address new researcher needs, such as a universal, robust object-level metadata system and additional visualization engines. Several other research groups have leveraged `bossDB` deployments, including NeuroData which serves approximately diverse collaborators in several new modalities (e.g., light microscopy, array tomography, Clarity) and added several new tools and capabilities to the `bossDB` ecosystem.

One concern about running a cloud data archive is estimating and managing cost. `bossDB` architecture was designed to allow dynamic scaling of resources to balance cost with performance and throughput capacity. As our software stack continues to mature, we plan to further optimize our tiered storage architecture (e.g., automatic migration data between S3 Standard, Infrequent Access, and Glacier tiers). The proposed system will provide a framework that is able to trivially scale from terabytes to petabytes while maintaining a balance between cost efficiency and performance.

As modern neuroscience datasets continue to grow in size, the community is fortunate to have several options to store and share their data. The precompute format [18] offers a flexible, lightweight option that is readily deployable in both local and cloud settings. As mentioned above, DVID [11] is used to manage immutable and versioned annotations at the terascale level. We believe that our `bossDB` solution offers key advantages in scalability (adaptable from gigabyte to petabyte storage); authentication to manage user access workloads and costs; indexing to promote data exploration and discovery; and managed services to ensure that data is maintained and available in an efficient manner for a variety of user workflows. For a given research lab (or even within the lifecycle of a scientific question), one or more of these storage solutions may be most appropriate to enable and share results.

The standardization and scalability provided by our data archive will support a fundamental change in how researchers design and execute their experiments, and will rapidly accelerate the processing and reuse of high-quality neuroscience, most immediately for the large, petascale image and annotation volumes produced by IARPA MICrONS. No previously existing platform met the operational and scaling requirements of the program, including managing an estimated 3-5 petabytes of image and annotation data – much larger than public neuroanatomical data archives. The `bossDB` software and documentation is open source and we are eager to expand the user community, supported modalities, and features. More information, examples and support are available at bossdb.org and https://github.com/jhuapl-boss/.

## 5 Acknowledgements

We would like to gratefully acknowledge our collaborators at NeuroData, including Alex Baden, Kunal Lillaney, Randal Burns, Joshua Vogelstein, Ben Falk, and Eric Perlman; Daniel Xenes at JHU/APL; Priya Manavalan, Jacob Vogelstein, and Denise D’Angelo; and our user community.

This material is based upon work supported by the National Institutes of Health (NIH) grant R24MH114785 under the Data Archives Program, and by the Office of the Director of National Intelligence (ODNI), Intelligent Advanced Research Projects Activity (IARPA), via IARPA Contract No. 2017-17032700004-005 under the MICrONS program. The views and conclusions contained herein are those of the authors and should not be interpreted as necessarily representing the official policies or endorsements, either expressed or implied, of the NIH, ODNI, IARPA, or the U.S. Government. The U.S. Government is authorized to reproduce and distribute reprints for Governmental purposes notwithstanding any copyright annotation therein.

